# Downfield Proton MRSI with whole-brain coverage at 3T

**DOI:** 10.1101/2023.01.27.525726

**Authors:** İpek Özdemir, Sandeep Ganji, B.S. Joseph Gillen, Semra Etyemez, Michal Považan, Peter B. Barker

## Abstract

**Purpose:** To develop a 3D downfield magnetic resonance spectroscopic imaging (DF-MRSI) protocol with whole brain coverage and post-processing pipeline for creation of metabolite maps.

**Methods:** A 3D, circularly phase-encoded version of the previously developed 2D DF-MRSI sequence with 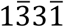 spectral-spatial excitation and frequency selective refocusing was implemented and tested in 5 healthy volunteers at 3T. Downfield metabolite maps with a nominal spatial resolution of 0.7 cm^3^ were recorded in 8 slices at 3T in a scan time of 22m 40s. An MRSI post-processing pipeline was developed to create DF metabolite maps. Metabolite concentrations and uncertainty estimates were compared between region differences for nine downfield peaks.

**Results:** LCModel analysis showed CRLB average values of 3-4% for protein amide resonances in the three selected regions (anterior cingulate (ACC), dorsolateral prefrontal cortex (DLPFC), and centrum semiovale (CSO)); CRLBs were somewhat higher for individual peaks but for the most part were less than 20%. While DF concentration maps were visually quite homogeneous throughout the brain, general linear regression analysis corrected for multiple comparisons found significant differences between CSO and DLPFC for peaks at 7.09 ppm (p= 0.014), 7.90 ppm (p=0.009), 8.18 ppm (p=0.009), combined amides (p=0.009), and between ACC and DLPFC for the 7.30 ppm peak (p=0.020). CRLB values were not significantly different between brain regions for any of the DF peaks.

**Conclusion:** 3D DF-MRSI of the human brain at 3T with wide spatial coverage for the mapping of exchangeable amide and other resonances is feasible at a nominal spatial resolution of 0.7 cm^3^.

## Introduction

In proton magnetic resonance spectroscopy (MRS) of the human brain, signals occur both upfield (UF) and downfield (DF) from the water resonance (1). A variety of compounds with aromatic or exchangeable (e.g. amine, amide, hydroxyl) groups resonate in the DF region (2). The amide protons of mobile proteins and N-acetylaspartate (NAA) are usually the most prominent DF signals in brain, but signals from ATP, histidine, homocarnosine, phenylalanine, glucose, glutathione, and nicotinamide adenine dinucleotide (NAD) are also known to be present and resonate in the DF region of the proton spectrum (3). Recently, small signals from I-tryptophan, a precursor of NAD(+) and serotonin syntheses, were also observed in DF MRS (4).

Although there have been comparatively few prior research studies using DF MRS, there are some indications of potential in either clinical or preclinical applications. For instance, DF-MRS has shown metabolic changes compared to normal brain in a murine brain glioma model (5), and changes in phenylalanine levels measured in patients with phenylketonuria (6). Also, estimations of brain pH have been shown to be feasible using the pH-dependence of the imidazole group resonances of histidine, with histidine levels increased using oral histidine supplementation (7).

Optimized detection of DF resonances depends on several factors: short echo time (TE) is important for minimization of T_2_ losses, since DF peaks typically have short T_2_ relaxation times (8) due to the effects of chemical exchange. In addition, since DF T_1_ relaxation times are also quite short (9), sensitivity can be enhanced by minimizing repetition times (TR) and also making use of the ‘relaxation enhancement’ effect due to chemical exchange (10). A factor of critical importance is to avoid saturation of the water signal (e.g. as is commonly used in UF MRS for water suppression) so that the DF resonances are not saturated via exchange; one way of avoiding water pre-saturation while still removing the water signal from the final spectrum is to use the metabolite cycling methods (11,12). However, other techniques are possible including spectral-selective excitation (13,14).

Most prior DF-MRS studies have used single voxel (SV) spatial localization (2,3,8,9,15–17). While SV MRS is usually rapid and generates high quality spectra, it does become prohibitively time-consuming and inefficient when multiple brain regions need to be measured. Recently, a single slice 2D MR spectroscopic imaging (MRSI) study of the DF resonances in normal human brain was published at a nominal spatial resolution of 1.5 cm^3^ using a 3T magnet (18). In this sequence, spectral-spatial excitation and frequency-selective 180° pulses were used to excite and refocus DF signals while avoiding saturation of the brain water magnetization. A relatively short TR was also used to maximize sensitivity via the ‘relaxation enhancement’ effect of exchange (or cross-relaxation) with the fully relaxed water signal (2,10,19,20). In the current communication, the further development of this sequence with increased spatial resolution and full brain coverage through the use of 3D-encoding is described. Results from 5 healthy volunteers are presented.

## Methods

Five healthy volunteers (2 females, age 32.6±13.7 years, max 57, min 25) were scanned using a 45-minute MR protocol using a Philips 3T ‘Ingenia Elition’ scanner equipped with a 32-receive channel head coil. All participants provided written informed consent approved by the Johns Hopkins Medicine Institutional Review Board (JHMIRB). Anatomical MRI, 3D downfield and 3D water reference MRSI scans were performed.

Prior to MRSI, B_0_ field homogeneity was optimized to 2^nd^ order using the a ‘FastMAP’ based technique (21). A 3D, circularly phase-encoded version of the previously developed 2D DF-MRSI sequence (18) with 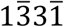 spectral-spatial excitation and frequency selective refocusing was implemented (Figure 1). All MRSI scans were performed with a FOV of 200×180×120 mm and a matrix size of 30×27×8 (elliptical k-space sampling), giving a nominal voxel size of 7×7×15 mm (≈ 0.7 cm^3^). Scan parameters were TR 287 ms, TE 22 ms, flip angle 78°, 1 excitation, scan time 22m 40s. The 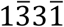 pulse was implemented with a maximum B_1_ of 22 μT and 5-lobe sinc pulses for slice selection, slice thickness 12 cm, and the delay (δ) set to give maximum excitation at 7.4 ppm. A frequency-selective sinc-Gauss 180° pulse applied at 7.8 ppm (11ms, 400 Hz BW) was used to refocus the downfield resonances. An inferior saturation pulse was also applied. Using an oblique axial prescription, spatial coverage was achieved from the base of the cerebellum to the vertex (Figure 2).

**Fig.1.**
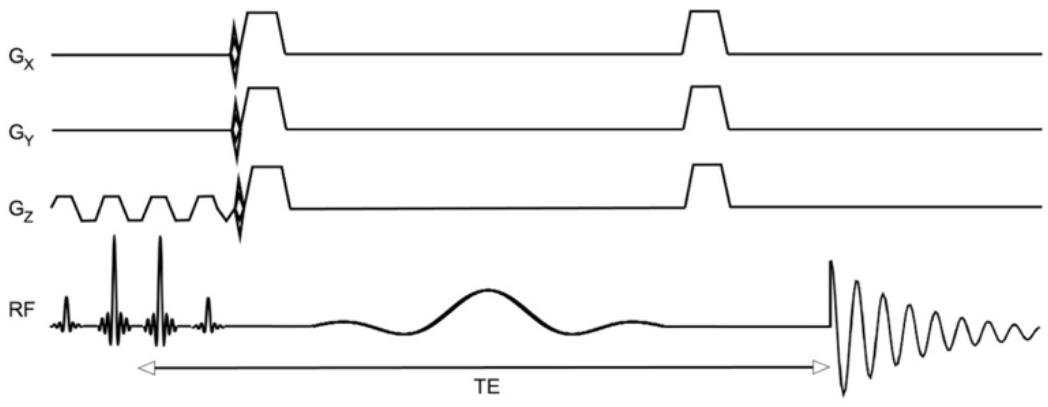
Pulse sequence for 3D-DF-MRSI. After 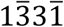 spectral-spatial excitation, phase-encoding is applied in 3 dimensions followed by frequency selective refocusing and acquisition of the spin echo signal.

**Fig.2.**
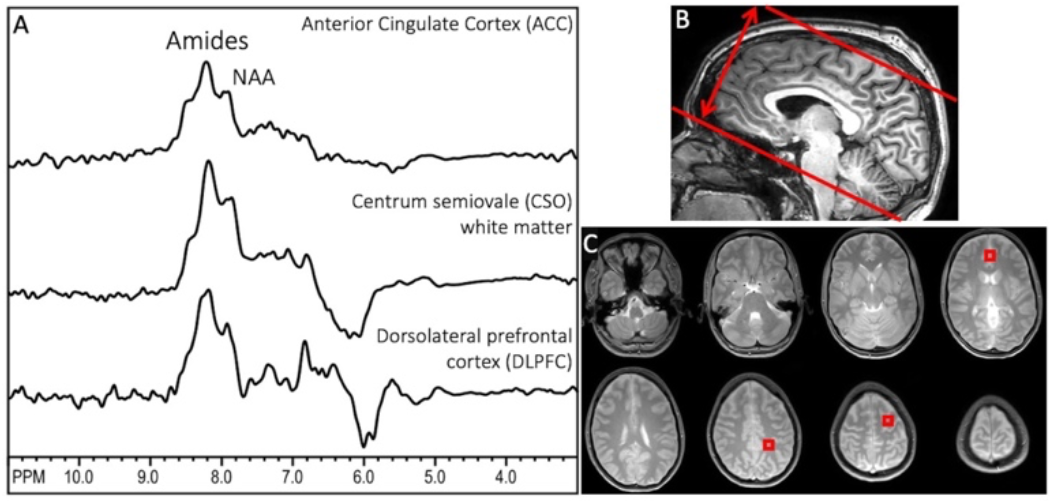
(A) Representative downfield spectra from one subject from the 3 brain regions used for quantitative analysis (anterior cingulate cortex (ACC), centrum semiovale (CSO), and dorsolateral prefrontal cortex (DLPFC), (B) sagittal T_1_-weighted MRI showing the oblique-axial slab prescription for 3D-DF-MRSI; the slab thickness is (double arrow) is 12 cm in this case, (C) proton-density MRI showing the 8 slice locations covered by 3D-DF-MRSI, including indicated voxel locations in ACC (slice 4), CSO (slice 6) and left DLPFC (slice 7). NAA and amide groups are the largest peaks at 7.9 and 8.2 ppm under the acquisition conditions used here.

A non-water suppressed FID-MRSI scan was also recorded at the same resolution and matrix size as the DF-MRSI. Scan parameters were TR 264 ms, TE 1 ms, flip angle 30°, 1 excitation, SENSE acceleration (R=2), scan time 11m 14s.

Reconstruction of downfield metabolite maps was performed using MATLAB code and LCModel fitting (22). DF-MRSI spectra were frequency corrected using information from the water reference scan prior to the LCModel fitting, and residual water was removed using an HLSVD filter (23–25). The basis set for spectral fitting consisted of nine individual Gaussian peaks as described previously (18), specifically 6.83, 7.09, 7.30, 7.48, 7.90, 8.18, 8.24, 8.37, 8.49 ppm. Baseline spline stiffness was controlled using the LCModel parameter “DKNTMN”=5 (18). A full list of LCModel control parameters is given in the supplemental information. DF peak areas from the LCModel, including the combined amide groups at 8.1-8.3 ppm, were expressed relative to the water signal in institutional units (‘i.u.’) for all spectra within the brain mask of the 3D MRSI-data. Brain masks were calculated from localized MRI scans using the ‘snakes’ active contour algorithm (37), an iterative region-growing image segmentation technique. Note that no attempts were made to estimate or adjust for regional variations in brain water content, or relaxation times. Metabolite maps were reconstructed in MATLAB using the concentration values from LCModel (estimated with a 8 Hz Lorentzian line broadening) and interpolated linearly by a factor of 8.

Maps of the distribution of the Bo field were reconstructed in one subject using the frequency of the water peak in the H_2_O-MRSI scan; in addition, the change in resonance frequency during the scan (i.e., frequency drift) due to changes in gradient temperature was estimated from the frequency of the water peak in the raw k-space data.

Metabolite levels and Cramer Rao Lower Bounds (CRLBs) of the DF signals were compared between three selected regions of interest in gray matter (anterior cingulate cortex (ACC), dorsolateral prefrontal cortex (DLPFC)), and white matter (centrum semiovale (CSO)). Up to four voxels were averaged for each brain region in each subject. R 3.5.1 was used to perform statistical analysis (36). After log transformation, general linear regression method was performed to compare brain metabolite levels between ACC, CSO, and DLPFC. The Benjamini-Hochberg (BH) procedure was used for multiple comparison correction. P values corrected with the BH procedure was considered significant if its value was smaller than 0.05.

## Results

Figure 2 shows representative spectra from the three voxels in different brain regions selected for analysis in one subject, as well as the slab thickness of 120 mm and proton density localizer images for the 8 slices covered by MRSI. NAA and amide groups are the largest peaks at 7.9 and 8.1-8.3 ppm.

Figure 3A shows an example LCModel fit from an ACC spectrum, while 3B and 3C show LCModel fits for the DLPFC and CSO regions.

**Fig.3.**
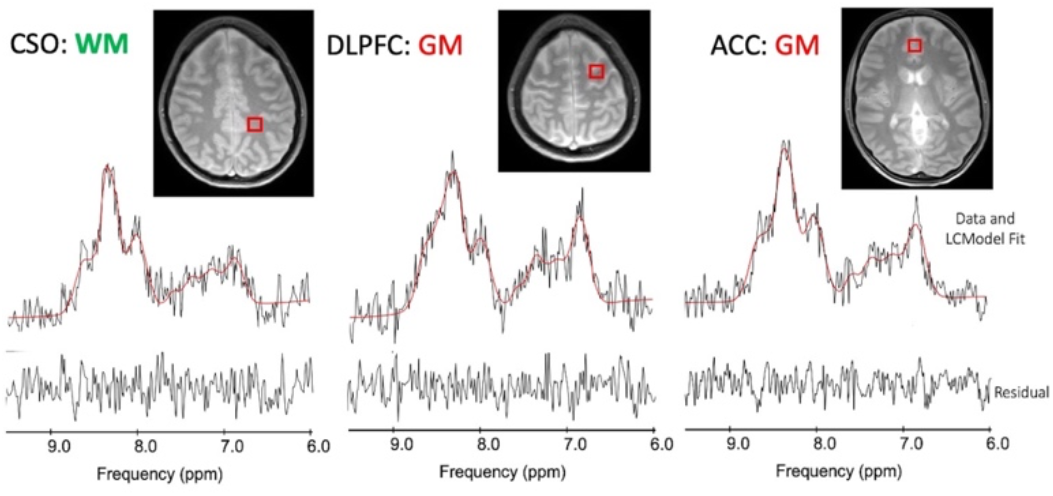
(A) Example LCModel outputs from the white matter left superior centrum semiovale (CSO) voxel in one subject, as well as gray matter voxels in (B) the left dorsolateral prefrontal cortex (DLPFC) and (C) mesial anterior cingulate cortex (ACC).

Figure 4 shows the results of the LCModel analysis (concentration levels and CRLBs) for the individual DF peaks, as well as the combined amide proton peaks, in all 5 subjects. For the combined amide resonances, concentration levels were in the range 5 to 8 (‘institutional units, ‘i.u.’) and CRLBs were on the order of 5%. The 7.90 ppm NAA amide peak concentration was around 2 i.u., with CRLB values in the 7-8% range. Individual, smaller peaks (including those with overlap with other resonances) had proton concentrations in the 1 to 2 i.u. range and somewhat larger CRLBs, on the order of 20% in some cases. For the amide concentration estimates, ACC was not significantly different from either CSO or DLPFC, while CSO and DLPFC were significantly different (p=0.009). There was no significant difference in CRLB values between ACC, CSO and DLPFC for any of the individual peaks, or the combined amide group. For other DF peaks, significant differences were found for metabolite concentrations in CSO and DLPFC for the metabolites at peaks 7.09 ppm (p= 0.014), 7.90 ppm (p=0.009), 8.18 ppm (p=0.009), and in ACC and DLPFC for metabolite 7.30 ppm (p=0.020).

**Fig.4.**
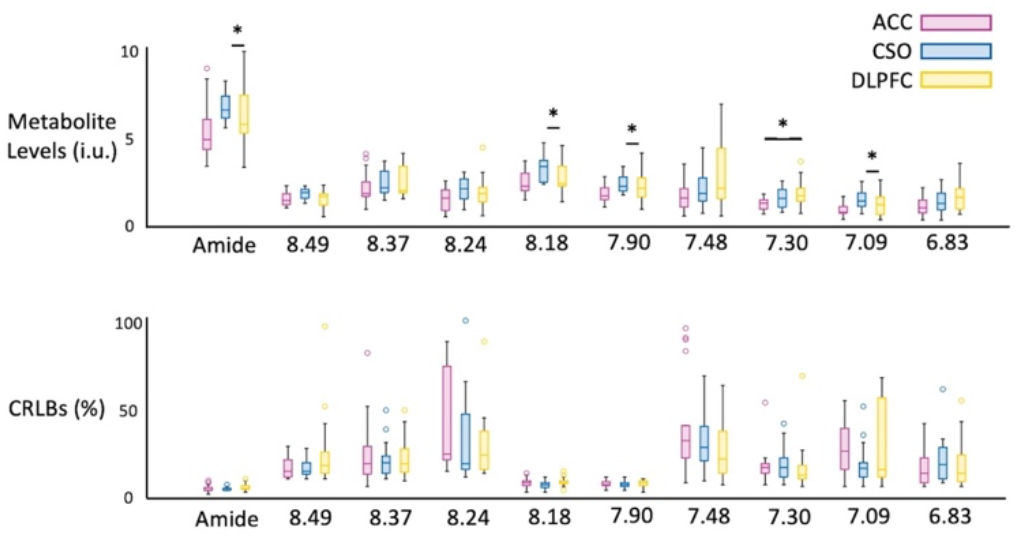
Descriptive statistics for (A) concentration and (B) CRLB values for the nine downfield peaks and combined amide resonances (8.1-8.3 ppm) in all 5 subjects. Significant differences were found estimated concentrations in CSO and DLPFC for peaks at 7.09 ppm (p= 0.014), 7.90 ppm (p=0.009), 8.18 ppm (p=0.009), and in ACC and DLPFC for metabolite 7.30 ppm (p=0.020). CRLB values were not significantly different between brain regions for any of the metabolites.

Figure 5 shows the reconstructed maps of the 7.90 ppm and combined amide resonances from all 8 slices in one subject. Relatively little contrast is seen between gray and white matter, but the spatial resolution is high enough to depict the ventricular CSF spaces with low signal. Loss of signal intensity in the most inferior and superior slices is due to the excitation profile of the slice (slab) selective 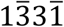 pulse. Reconstructed maps of the other DF peaks are given in the supplemental figure (S1).

**Fig. 5.**
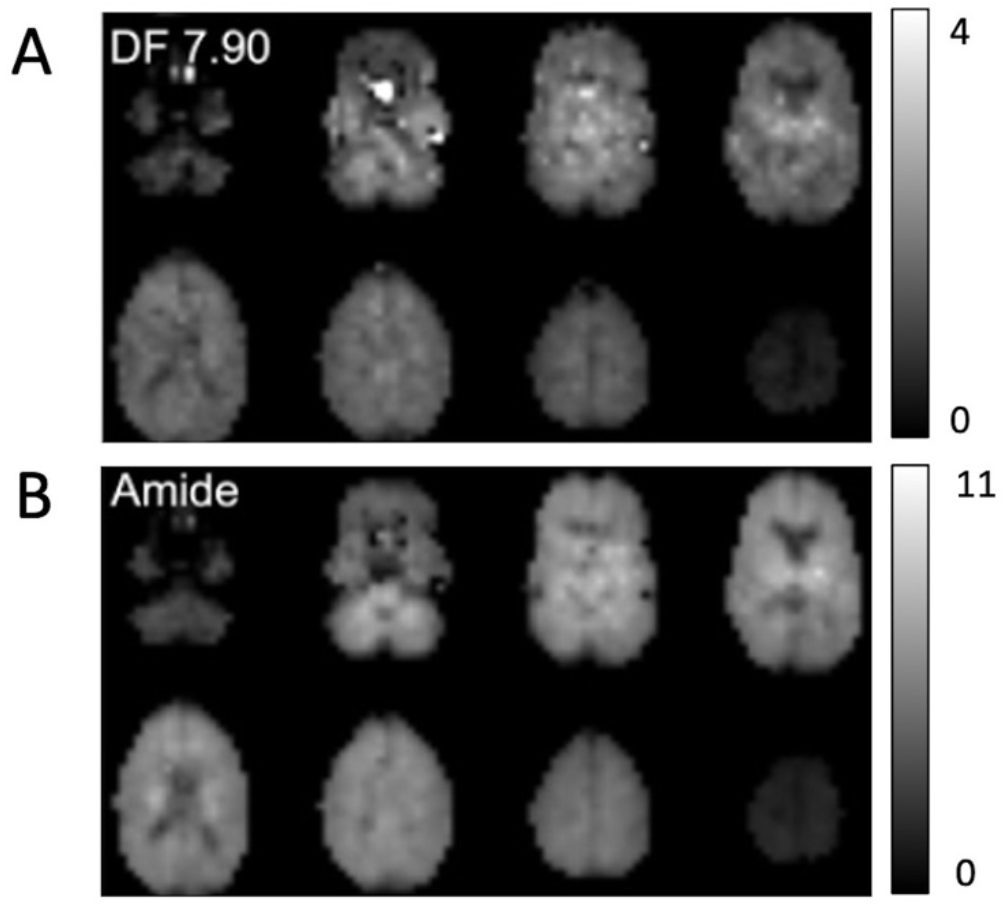
Example DF concentration maps from all 8 slices in 1 subject; maps are shown for (A) the NAA peak at 7.90 ppm as well as (B) the combined amide resonances (8.1-8.3 ppm). The maps are relatively homogeneous and show little gray-white contrast; however spatial resolution is high enough to depict CSF spaces such as the lateral ventricles which have low signal. There is some artifactual hyperintensity in the lower NAA slices attributable to residual water excited because of magnetic susceptibility effects (i.e. frequency shifts) due to adjacent air spaces.

Figure 6 illustrates B_0_ field inhomogeneity data, both as frequency shift maps from the H_2_O-MRSI scan as well as in histogram form, and, in addition, shows the frequency drift over the time course of the 11-minute H_2_O-MRSI scan.

**Fig.6.**
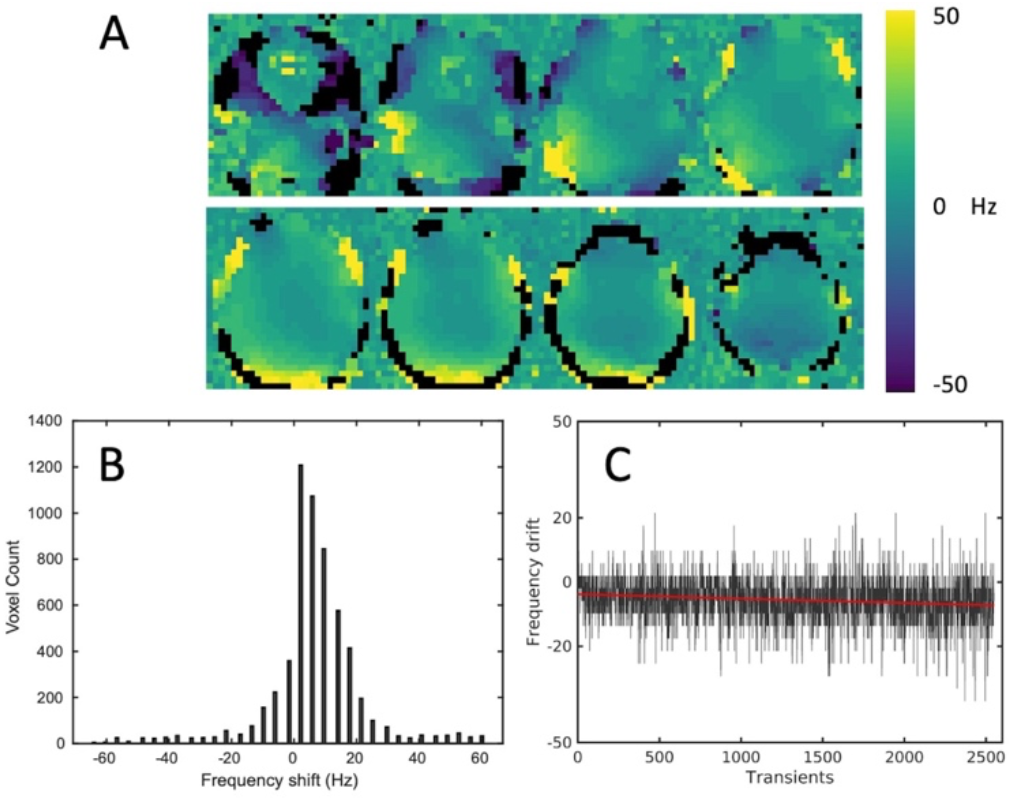
B_0_ field distribution over the 8 slices used for DF-MRSI in one subject: (A) B_0_ field maps in each slice (color map +/− 50 Hz), (B) histogram of frequency shift values within the brain over all 8 slices, (C) frequency drift in Hz measured from the water MRSI scan of 11 minutes duration (approximately 2500 transients in total). It can be seen that both field homogeneity over the volume of the brain, as well as Bo stability, are good in this subject.

## Discussion

This study shows that downfield 3D-MRSI with near whole brain coverage and ≈ 1 cm^3^ spatial resolution is possible at 3T in around a 20-minute scan time. Compared to upfield MRSI, DF-MRSI has some technical advantages, including the ability to use short TR (due to the short T_1_s and ‘relaxation enhancement’ effect), no need for additional water or lipid suppression, and slightly less sensitivity to field inhomogeneity, since the downfield resonances have broader intrinsic linewidths.

T_1_ relaxation times and exchange rates of downfield peaks from, 6.84 to 8.50 ppm have previously been measured in the human brain at 3T (9). T_1_’s varied from as short as 176 to 525 ms, appreciably shorter than those of upfield resonances which are typically more than 1 s (35). In addition, exchange rates (k_ex_) varied from 0.5 to 8.9 Hz; when the water signal is fully relaxed, the apparent relaxation rate (R_1a_ = 1/T_1a_) of exchanging peaks is given by R_1a_ = R_1a_ + k_ex_ (1). For the 8.24 ppm amide resonance, with a T_1_ of 254 ms and k_ex_ of 7.5 Hz, the apparent T_1_ (T_1a_) is calculated to be as short as 87 ms; thus with the TR in the current experiments of 287ms, the Ernst angle (26) for optimum sensitivity is very close to 90° (87.8°). Other DF peaks have different T_1_’s and exchange rates; for instance, the 7.86 ppm amide peak of NAA has a longer T_1_ (525 ms) and slower exchange rate (2.4 Hz), so a much longer apparent T_1_ (415 ms). Therefore, not all peaks benefit equally from the relaxation enhancement effect, and to optimize sensitivity for the NAA amide peak would require a lower flip angle (Ernst angle = 59.9° for TR 287 ms). Similarly, the frequency offset for maximal excitation of the 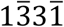 excitation pulse (7.4 ppm in the current experiments) and as well as the bandwidth and frequency offset of the refocusing (400 Hz/7.8 ppm) can be adjusted for preferential detection of different DF peaks. The current choice of parameters reflects a desire to maximize signals between 6 to 9 ppm with an emphasis on amide resonances; the 180° pulse refocusing profile also plays a role in quality of water suppression, since the 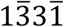 pulse alone does generate an appreciable residual water *in vivo*.

Another factor to consider for 3D acquisitions with extended brain coverage is how Bo field inhomogeneity over the brain may affect both detection efficiency and the presence of artifacts. Using 2^nd^ order shimming methods, field mapping experiments (Figure 6) indicate that over 90% of brain voxels have frequency offsets due to Bo inhomogeneity of less than +/− 15 Hz, which should have negligible influence on MRSI results. However, voxels adjacent to the air spaces of the paranasal sinuses and external auditory canal may show shifts of as much as ~50 Hz, which can result in substantial excitation and refocusing of the water signal in these regions. These are regions where typically upfield MRS or MRSI is also not possible due to Bo inhomogeneity (27). In addition to Bo inhomogeneity, the influence of Bo drift also needs to be considered in the quite lengthy 3D MRSI scans used here. Again, the drift measurements shown in figure 6 show that (at least for the 3T system and pulse sequence used in the current experiments), the effects of field drift are negligible (< 5 Hz over an 11-minute scan duration).

Inspection of DF maps also indicates reduction of signal intensity in the most inferior and superior slice locations, due to the excitation profile of the slice selection RF pulses used in the 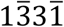 pulse. The rectangularity of the slice profile was improved by adding additional side-lobes of the sinc pulses compared to those used in the prior 2D sequence; while this improves the slice profile, it also limited the maximum flip angle to 78° with the timing parameters used and a maximum B_1_ level of 22 μT. The high B_1_ level used resulted in high bandwidth for the individual elements of the 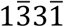 pulse (≈ 11.6 kHz), so that the 3D slab excited experienced minimal chemical shift displacement effects (CSDE). For the frequency range of peaks analyzed in these scans (6.8 to 8.5 ppm, i.e. a frequency range of 1.7 ppm (~218 Hz)), the CSDE corresponded to approximately 2.3 mm only.

Although the current scan time for DF-MRSI was 22 minutes, it should be relatively straight forward to decrease this to under 10 minutes through further sequence optimization, and the use of fast-MRSI techniques, for instance, parallel-imaging approaches (e.g. SENSE-(28) and GRAPPA-MRSI (29)), sparse sampling (30), or readout gradients during data acquisition such as echo-planar spectroscopic imaging (EPSI) (31,32) or concentric circles (33).

Finally, amide proton transfer (APT) chemical exchange saturation transfer (CEST) MRI has been shown to have value in evaluating patients with brain tumors (34); DF-MRSI may give specific metabolic information on slowly exchanging molecules that is complementary to that observed using APT-CEST MRI, which is more sensitive for intermediate exchange rates. Compared to APT-CEST, DF-MRSI offers somewhat greater spectral resolution but lower spatial resolution, so may offer additional metabolic information beyond that available from CEST.

In summary, 3D downfield MRSI of the human brain for the mapping of exchangeable amide and other resonances is feasible at 3T at a nominal spatial resolution of 0.7 cm^3^, and may be of use in future studies of brain tumors or other neuropathological conditions.

## Acknowledgements

Supported in part by NIH grants R01EB028259 and P41EB031771.

**Supporting Information Figure S1 -** Reconstructed estimated concentrations maps (i.u.) for DF peaks (A) 8.49 ppm, (B) 8.37, 8.24 and 8.18 ppm peaks (amides), (C) 7.48, 7.30 and 7.09 ppm, and (D) 6.83 ppm. The maps are generally quite homogeneous, but for the smaller and overlapping peaks exhibit several artifacts which perhaps could be removed with more advanced post-processing techniques. For instance, hypointense pixels in the 8.24 ppm map seem to correspond to hyperintense pixels in the 8.18 ppm, likely because of incorrect fitting of overlapping peaks.

**<u>Supplemental Information Table 1 - *Example Control file (.control) for LCModel analysis*</u>**

